# Production of very-high-amylose cassava by post-transcriptional silencing of branching enzyme genes

**DOI:** 10.1101/477414

**Authors:** Wenzhi Zhou, Shanshan Zhao, Shutao He, Qiuxiang Ma, Xinlu Lu, Xiaomeng Hao, Hongxia Wang, Jun Yang, Peng Zhang

**Author notes:** Equal contributions (Wenzhi Zhou, Shanshan Zhao). Correspondence should be addressed to P. Z.

## Abstract

High amylose starch, a desired raw material in the starch industry, can be produced by plants deficient in the function of branching enzymes (BEs). Here we report the production of transgenic cassava plants with starches containing up to 50% amylose due to the constitutive expression of hair-pin dsRNAs targeting the BE1 or BE2 genes. A significant decrease in BE transcripts was confirmed in these transgenic plants by quantitative real-time RT-PCR. The absence of BE1 protein in the BE1-RNAi plant lines (BE1i) and a dramatically lower level of BE2 protein in the BE2-RNAi plant lines (BE2i) were further confirmed by Western blot assays. All transgenic plant lines were grown up in the field, but with reduced biomass production of the above-ground parts and storage roots compared to wild type (WT). Considerably high amylose content in the storage roots of BE2i plant lines was achieved, though not in BE1i plant lines. Storage starch granules of BE1i and BE2i plants had similar morphology as WT, however, the size of BE1i starch granules were bigger than that of WT. Comparisons of amylograms and thermograms of all three sources of storage starches revealed dramatic changes to the pasting properties and a higher melting temperature for BE2i starches. Glucan chain length distribution analysis showed a slight increase in chains of DP>36 in BE1i lines and a dramatic increase in glucan chains between DP 10-20 and DP>40 in BE2i lines, compared to that of WT starch. Furthermore, BE2i starches displayed a B-type X-ray diffraction pattern instead of the A-type pattern found in BE1i and WT starches. Therefore, cassava BE1 and BE2 function differently in storage root starch biosynthesis; silencing of cassava BE1 or BE2 caused various changes to starch physico-chemical properties and amylopectin structure. We also report that remarkably high amylose content in cassava starch has been first obtained in transgenic cassava by silencing of BE2 expression, thus showing a high potential for future industrial utilization.

## Introduction

Cassava (*Manihot esculenta* Crantz) is one of the most important starchy crops worldwide, providing staple food for more than 700 million people in the tropics and subtropics (Hershey, 2017). Compared with other starches (e.g., from maize and potato), pure-white cassava starch contains low levels of fat (0.08–1.54%), proteins (0.03–0.6%), and phosphorus (0.75–4%), making it an excellent starch resource for various industrial applications (Richard et al., 1991; Moorthy, 2002; Balagopalan, 2002; Sharma et al., 2016). Modified starches of cassava have been broadly used in the food, textile, pharmaceutical, paper manufacturing, and other industries (Zhu et al., 2015). Promotion of cassava-based bioethanol production in China and other South-east Asian countries also made the crop an important source of bioenergy development over the last decade (Marx, 2019).

To meet the growing industrial demands for value-added starches, the native starch with novel physico-chemical property is needed in cassava (Zhang et al., 2017). Unlike the traditional physical and/or chemical starch modifications, breeders intend to produce new cassava varieties with modified starches to expand their application spectrum through genetic approaches, such as starch-related mutant identification and reverse-genetic/biotechnological approaches (Schwall et al., 2000; Jobling et al., 2002; Raemakers et al., 2005; Ceballos et al., 2007; Zhao et al., 2011; Bull et al., 2018).

Therefore, the development of cassava varieties having novel starches including waxy (amylose-free) and high-amylose starches are important objectives for cassava breeders. The target genes are mainly involved in starch biosynthesis, such as granule bound starch synthases (GBSS), soluble starch synthases (SSS), starch branching enzyme (SBE or BE), and debranching enzyme (DBE) (Zeeman et al., 2010; Bahaji et al, 2014). More specifically, the SSS, BE, and DBE are involved in amylopectin synthesis, and GBSS is the key enzyme for amylose biosynthesis in plants including cassava (Zhao et al., 2011; Bull et al., 2018).

Since the BE gene was first studied by Mendel with the wrinkled (rugosus, r) phenotype in the pea (*Pisum sativum* L.) mutants (Bhattacharyya et al., 1990; Tetlow and Emes, 2014) and identified from maize and rice using the well-known *amylose-extender* (*ae*^-^) mutants (Hedman and Boyer, 1982; Mizuno et al., 1993; Kim et al., 1998), its important role in amylopectin biosynthesis has been studied in many plants like potato (Rydberg et al., 2001), maize (Yao et al., 2004), wheat (Regina et al., 2005), and rice (Nakamura et al., 2012). BE has the unique activity of producing α-1, 6 glycosidic bonds, the basis in forming branching chains of amylopectin. BEs usually comprise two classes, BE1 and BE2, which can be distinguished by their amino acid sequences (reviewed by Tetlow and Emes, 2014). Enzymatic activity of BEs is regulated by phosphorylation and protein-protein interactions (Tetlow et al., 2004). Suppression of BE function results in the increase of amylose content in many plants, for example, potato (Schwall et al., 2000), wheat (Sestili et al., 2010), rice (Sun et al., 2017), and sweet potato (Shimada et al., 2006). Normally, high-amylose starches show altered properties in applications distinguishable from the native or waxy starches, including their delayed gelatinization temperature, high gelling capacity, and easy forming film (Richardson et al., 2000; Zhou et al., 2015; Wang et al., 2017). In cassava, studies related to cloning and expression analysis of BE genes have been previously reported (Salehuzzaman et al., 1992; Baguma et al., 2003; Pei et al., 2015). With the recently release of the cassava genome sequence (Prochnik et al., 2012; Wang et al., 2014; Bredeson et al., 2016) and development of cassava transformation technology (Liu et al., 2011; Zhang et al., 2017), we now have the capacity to breed novel cassava cultivars with desirable traits using state-of-the-art technologies including CRISPR/Cas9-based genome editing (Odipio et al., 2017; Sun et al., 2017; Bull et al., 2018; Gomez et al., 2018).

Amylose content of native cassava starch is about 21% on average, with a range from 15%-27% (Sánchez et al., 2009). Although cassava mutants having amylose-free or small-granule starch with relatively higher amylose (30%) have been identified from self-pollinated generations in germplasm collections at the International Center for Tropical Agriculture (CIAT, Ceballos et al., 2007, 2008), there are still no reports on very-high-amylose cassava. Due to the difficulty and time-consuming processes needed to generate high amylose cassava cultivars via traditional breeding programs, transgenic approaches became an alternative means of achieving this goal, and have proven very effective for the rapid acquisition of cassava lines with novel starches (Raemakers et al., 2005; Ihemere et al., 2006; Zhao et al., 2011; Radhika et al., 2014). Furthermore, GBSS-RNAi transgenic cassava lines have generated many more diversified starches than the *gbss* mutant AM206-5 from CIAT (Zhao et al., 2011, Rolland-Sabaté et al., 2013), providing us with sound evidence that this approach will be effective.

In this study, we successfully developed transgenic cassava lines with high amylose starch in their storage roots by the silencing of BE1 and BE2 expression. The transgenic cassava shows altered phenotypes and properties of storage starch, indicating a different function for each of these genes in starch biosynthesis. Importantly, these high amylose starches showed B-type XRD diffraction-grams, with higher melting temperatures and potentially useful pasting properties, thus providing very promising raw materials for industrial applications.

## Materials and methods

### Production and growth of transgenic cassava

To repress the expression of starch branching enzymes BE1 and BE2, the binary vectors p35S::BE1-RNAi and p35S::BE2-RNAi were constructed using the pRNAi-dsAC1 plasmid backbone (Vanderschuren et al., 2009). The AC1 sequence was replaced with a partial cDNA sequence of *MeBE1* (GenBank accession No. MK086025, 2102-2299 bp) and a partial cDNA sequence of MeBE2 (Genbank accession No. MK086026, 1649-1947 bp) in p35S::BE1-RNAi and p35S::BE2-RNAi, respectively. The resulting constructs were introduced into *Agrobacterium tumefaciens* LBA4404 and the cassava cultivar TMS60444 was used to generate transgenic plants as described previously (Zhang et al., 2000a). Genomic integration of T-DNA in the transgenic cassava plants was confirmed using standard protocols for Southern blot analysis with DIG-labeled PCR products of the hygromycin phosphotransferase gene (*HPT*) for hybridization (Zhao et al., 2011).

The MeBE1 and MeBE2 RNAi transgenic lines (BE1i and BE2i for short, respectively) and wild type (WT) cassava TMS60444 were propagated by *in vitro* shoot culture followed by transfer to pots in the greenhouse for macropropagation (16 h/8 h of light/dark, 30 °C/22 °C day/night). In field trials, ten stem cuttings each per transgenic line and WT were planted in the field in early May at the Wushe Plantation for Transgenic Crops, Shanghai, China (31°13948.0099N, 121°28912.0099E). Phenotypic data on the performance of plants was recorded and the storage roots were harvested in early November for agronomic trait evaluation and subsequent experiments.

### Transcriptional and translational expression analysis of *MeBE1* and *MeBE2* genes

Leaves from three plants per line were harvested from 5-month-old plants and then ground in liquid nitrogen for mRNA extraction. To quantify the expression of *MeBE1* and *MeBE2* genes, real-time qRT-PCR was performed as described previously (Xu et al., 2013). The β-Actin was used as a reference for normalization. The primers were as follows:

*MeBE1* (forward, 5′-GCTCGCACTTGTGTGGTTTA-3′; reverse,

5′-CATCGGCAATCAAAGAAGGT-3′), MeBE2 (forward,

5′-CAGTTCAAGCACCAGGTGAA-3′; reverse,

5′-AAGCTTTTTGATGCGAGGAA-3′), and

β-Actin (forward, 5′-TGATGAGTCTGGTCCATCCA-3′; reverse,

5′-CCTCCTACGACCCAATCTCA-3′).

Total protein was extracted from cassava leaves as described by Ritte et al. (2000) and quantified according to the method of Bradford (Bradford, 1976). Approximately 60 μg of protein was loaded on an SDS-PAGE gel and Western blotting was performed with rabbit antisera against MeBE1 and MeBE2 proteins, respectively.

### Starch extraction and analyses of amylose content and starch granule size

The starch was extracted from storage roots that were harvested from six-month-old cassava plants grown in the field, washed twice in 75% ethanol, and air-dried overnight at 40°C (Zhao et al., 2011). The amylose content of starch was determined using a previously described colorimetric method (Knutson and Grove, 1994). Amylose type III and amylopectin (Sigma A0512 and Sigma 10118, St. Louis, MO, USA) from potato were used to establish the standard curve for amylose quantification.

To observe starch granules by scanning electron microscopy (SEM), isolated starch granules in distilled water were placed or sprinkled on double-sided sticky tape, air-dried, and coated with gold powder. Samples were observed and photographed by SEM (JSM6360lv, JEOL, Tokyo, Japan). The granule size distribution of starch was determined as described (Zhou et al., 2015). A Master-size 2000 laser diffraction instrument (Malvern Instruments Ltd., Worcestershire, UK) was used in wet-well mode. The starch was added to the reservoir and sonicated for 30 s at 6 W until an obscuration value of 12–17% was achieved. The refractive indices used for the water and starch were 1.330 and 1.50, respectively.

### Physicochemical property assay of cassava storage starches

The chain length distribution was determined following the protocol previously described by Ritte et al. (2000), using high performance anion exchange chromatography (HPAEC) that was equipped with a CarboPac PA1 column. Sample preparation included boiling 5 mg starch and enzymatic digestion with isoamylase (I5282, Sigma, St. Louis, MO, USA). Oligosaccharides with a polymerization degree of 4–7 (47265, Sigma, St. Louis, MO, USA) were used as a standard.

X-ray diffraction analysis of the extracted starches was performed using a D8 Advance Bruker X-ray diffractometer (Bruker AXS, Karlsruhe, Germany) and scanned through the 2θ range of 5–60° at a rate of 4° min^−1^. The starch pasting properties were analyzed using a rapid viscosity analyzer (model RVA-Super 4; Newport Scientific Pty. Ltd., Australia). The starch was suspended in distilled water (5% w/v, dry weight basis, 25 ml) and tested using a dedicated program for cassava. The temperature of the starch slurry was increased from 30°C to 95°C at a rate of 5°C min^−1^ and held at 95°C for 6 min, followed by cooling to 50°C at the same rate and maintenance for 10 min. The rotating speed of the paddle remained constant (160 rpm) throughout the analysis, excluding the speed of 960 rpm applied during the first 10 s. The thermal properties of the starch samples were analyzed using a differential scanning calorimeter (DSC, Q2000; TA Instruments Ltd., Crawley, UK) at a temperature scanning range of 30–95°C.

### Statistical analysis

Root samples were collected from three independent plants per line and then pooled for further analyses. Data from at least three replicates were presented as the mean ± SD. Analysis of variance (ANOVA) by Duncan’s multiple comparison tests or independent samples Student’s t-test was performed using SPSS software, version 17 (SPSS Inc., Chicago, IL, USA). An alpha value of P< 0.05 was considered statistically significant.

## Results

### Down-regulation of MeBE1 and MeBE2 expression in cassava leads to reduced growth of field-grown plants

Among transgenic cassava plant lines, 3 BE1i lines and 4 BE2i lines showing single T-DNA integration in their genome were selected for further evaluation. After 6 months of growth in the field, phenotypic measurements were performed on the WT and transgenic lines, including plant height, fresh-weight biomass, root number, root length, and root diameter (Fig.1). The average height was 2.2 m for WT. The heights ranged from 1.7 m (BE1i-13) to 2.1 m (BE1i-18) for BE1i lines, and 1.4 m (BE2i-24) to 2.0 m (BE2i-58) for BE2i lines (Fig 1 a, b), both of which were shorter than WT. The biomass per plant ranged from 0.7 kg (BE1i-13) to 1.6 kg (BE1i-26) for BE1i lines and 0.9 kg (BE2i-24) to 1.4 kg (BE2i-38, BE2i-59) for BE2i lines (Fig 1 a, c), which is much lighter than the average fresh weight of WT (3.5 kg). The average root number per plant reached from 10 (BE1i-13) to 16 (BE1i-18) for BE1i lines and 12 (BE2i-24, BE2i-59) to 14 (BE2i-58) for BE2i lines (Fig 1 a, d), which is dramatically less than of the WT average (24). The average root length was 20.8 cm for WT, and it ranged from 16.6 cm (BE1i-18) to19 cm (BE1i-13, BE1i-26) for BE1i lines and 14 cm (BE2i-24) to 21 cm (BE2i-59) for BE2i lines (Fig 1 a, e). The average root diameter was 2.8 cm for WT, and 2.3 cm (BE1i-18) to 2.6 cm (BE1i-13) for BE1i lines, and 2.3 cm (BE2i-58) to 2.5 cm (BE2i-24) for BE2i lines (Fig 1 a, f). Thus, for both BE1i and BE2i transgenic lines, the total biomass was reduced due to retarded growth of both aerial and subterranean organs.

**Figure 1.**
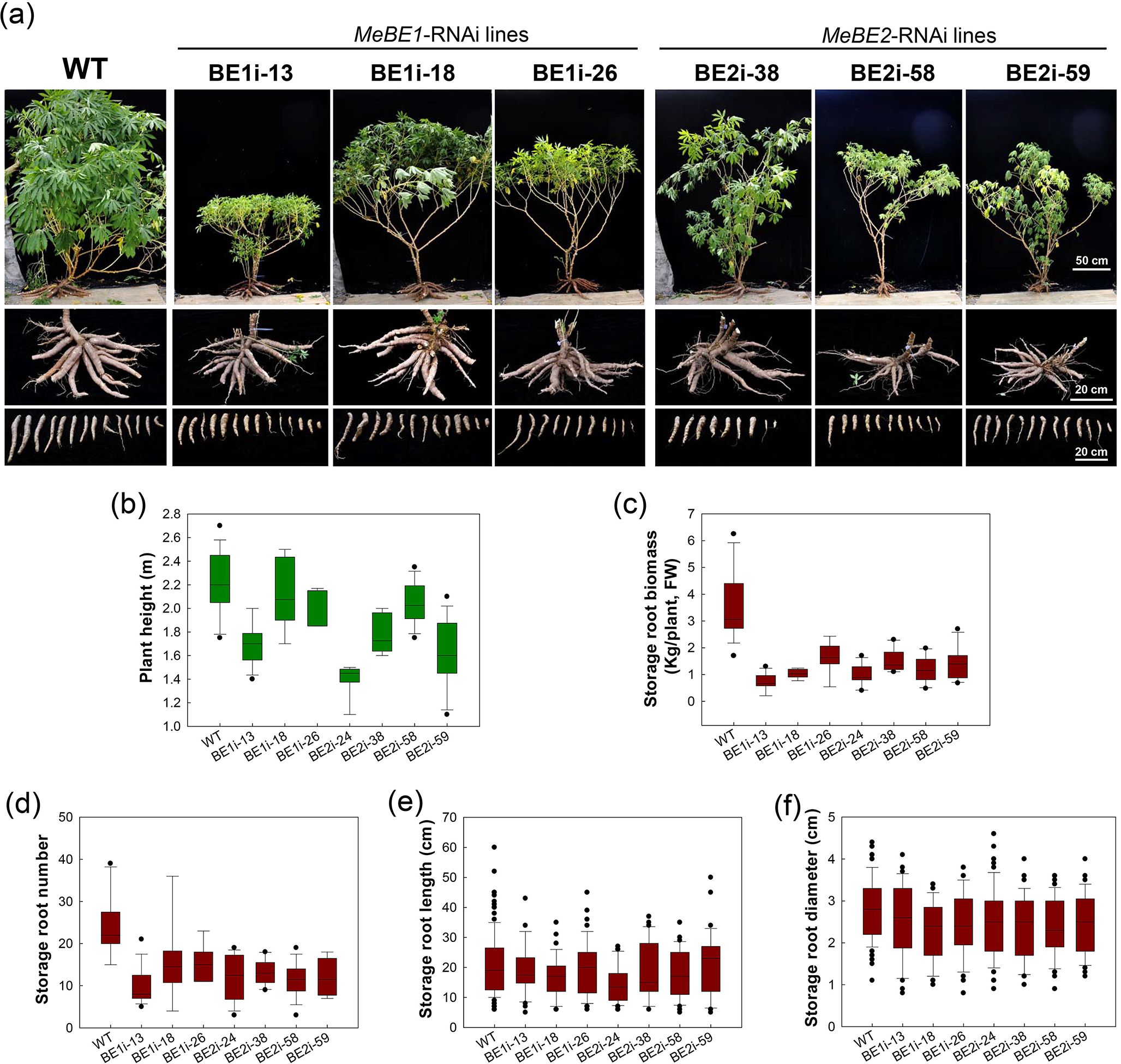
Phenotypes of field-grown cassava plants. (a) Canopy architecture (upper panel), attached storage roots (middle panel), and storage roots (lower panel) of the BE1i and BE2i transgenic plants in comparison with the wild type (WT). (b-f) Comparisons of plant height (b), storage root biomass (c), storage root number (d), storage root length (e), and storage root diameter (f).

To verify the phenotypic changes correspond with expression of BEs, transcriptional analysis by real-time qRT-PCR and translational analysis using Western blot assays were carried for MeBE1 and MeBE2 genes using leaf samples from field-grown cassava plants. Relative transcript levels of *BE1* and *BE2* genes were drastically reduced by 10-fold compared to WT in the BE1i or BE2i lines, respectively (Fig. 2a, b). No obvious change was detected for MeBE1 in BE2i lines and MeBE2 in BE1i lines (Fig2a, b). Furthermore, the protein bands of MeBE1 were undetectable in BE1i lines, while the MeBE2 bands in BE2i lines similarly showed a dramatic reduction (Fig. 2c). The protein levels of MeBE1 in BE2i lines and MeBE2 in BE1i lines did not change (Fig2c). These results suggested that the expression levels of BE1 and BE2 genes in cassava were specifically and individually silenced in the BE1i and BE2i transgenic lines, respectively.

**Figure 2.**
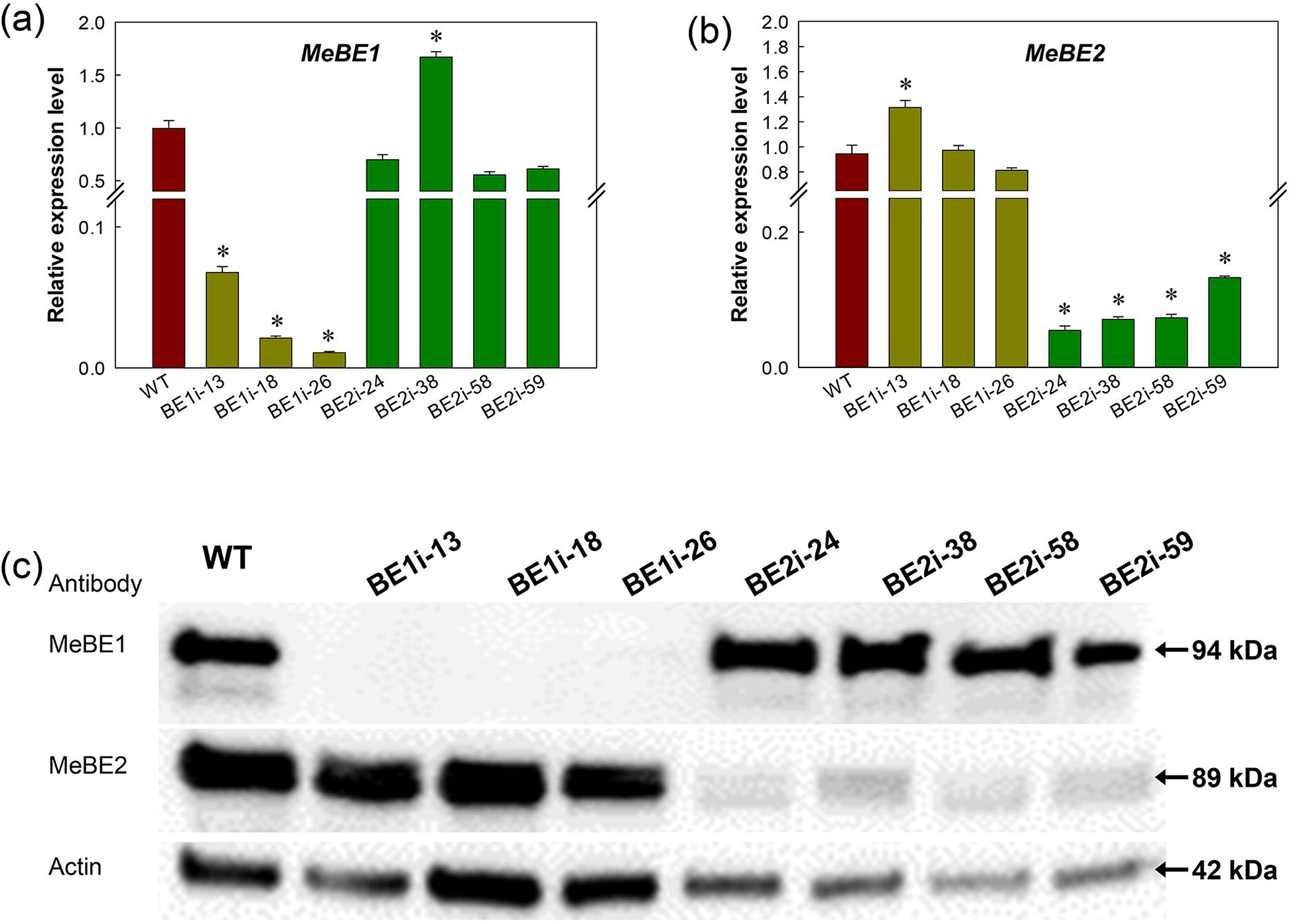
Expression of BE genes in cassava plants. The transcription (a) and translation (b) levels of BE1 and BE2 genes was drastically reduced in leaves of BE1i and BE2i lines, respectively. WT, wild type; BE1i-x, BE1i transgenic plant lines; BE2i-x, BE2i transgenic plant lines. Actin protein was used as a loading control in Western analysis. Significance was determined by the Student’s *t*-test at * *P* < 0.05.

### MeBE2 but not MeBE1 influences the amylose content in storage roots of cassava

Using colorimetric measurement of amylose-binding stain as a proxy for starch content, 22% amylose was detected in storage starch of WT. Amylose content in BE1i was almost the same as in WT (Fig. 3). However, for BE2i lines, the percentage of amylose in total starch was 50% in BE2i-24, 46% in BE2i-58 and 31% in BE2i-59, a dramatic increase over WT. These data support the hypothesis that BE2 plays key role in amylopectin biosynthesis in cassava storage roots, and that down-regulation of *MeBE2* expression by RNAi is an effective way of producing high-amylose starch in cassava.

**Figure 3.**
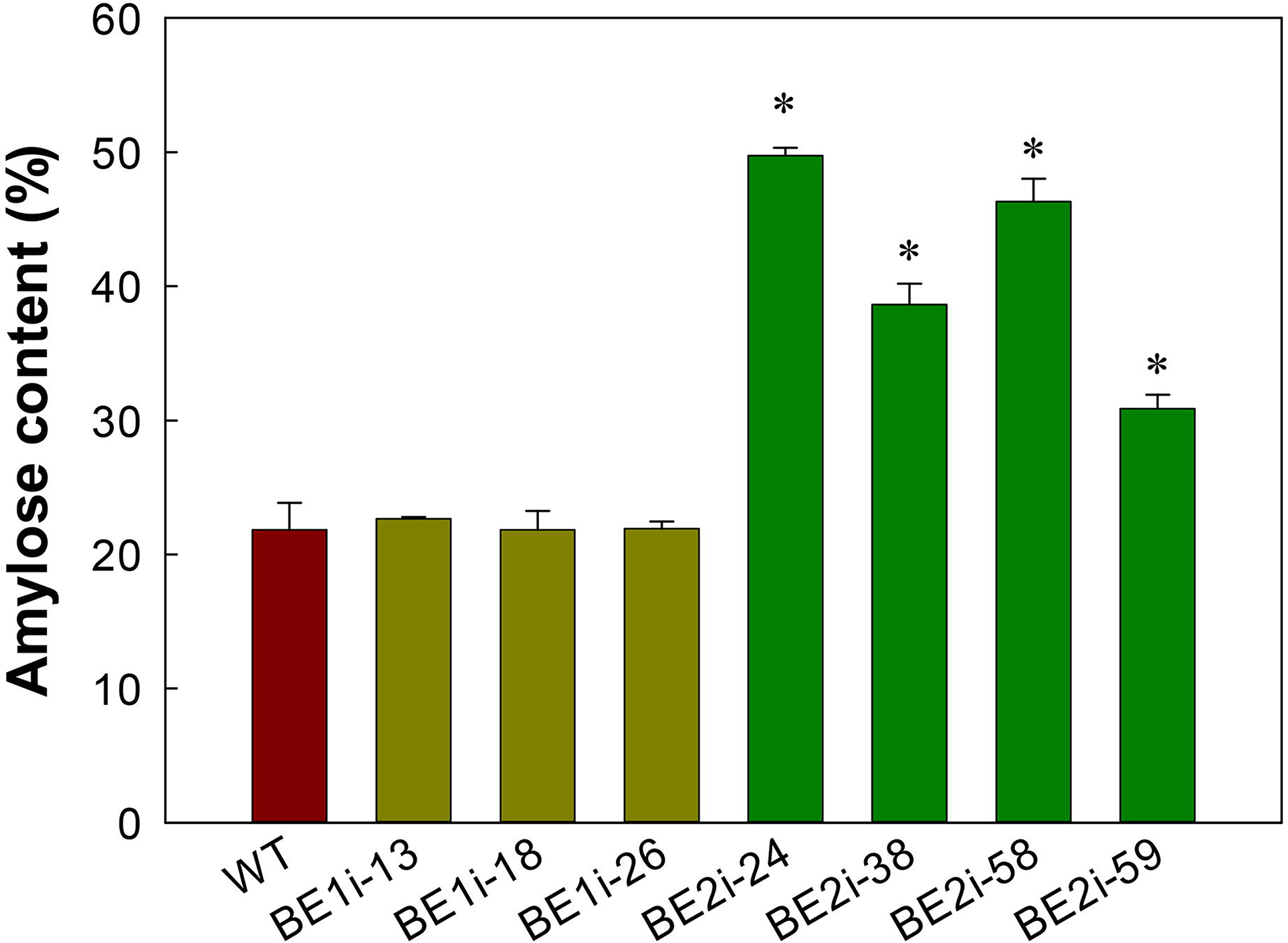
Amylose content of extracted starches from cassava storage roots of wild-type (WT) and BE-RNAi transgenic plants. BE1i-x, BE1i transgenic plant lines; BE2i-x, BE2i transgenic plant lines. Significance was determined by the Student’s *t*-test at * *P* < 0.05.

### MeBE1 predominantly affects starch granule size in cassava storage roots

Observation by scanning electron microscopy (SEM) revealed no clear difference in the starch granule shape among WT and BE1i or BE2i transgenic lines. All of these starch granules were a mixture of round, truncated, and dome-shaped granules (Fig. 4a). The diameter of starch granules from the WT and transgenic lines was compared using representative diameters Dx10, Dx50, and Dx90. In all BE1i lines, a significant increase in these values was detected, about 8 μm, 14 μm, and 22 μm, compared to 7 μm, 12 μm and 19 μm in WT starch. For BE2i lines, only BE2i-24 showed increased values of Dx10, Dx50 and Dx90 at about 9 μm, 15 μm and 25 μm, respectively. No significant changes for BE2i-38 or BE2i-59 starches were detected when compared to starch in WT (Fig4 b). In addition, the Dx90 of BE2i-58 starch granules was 21 μm, slightly larger than WT, but still smaller than BE1i lines.

**Figure 4.**
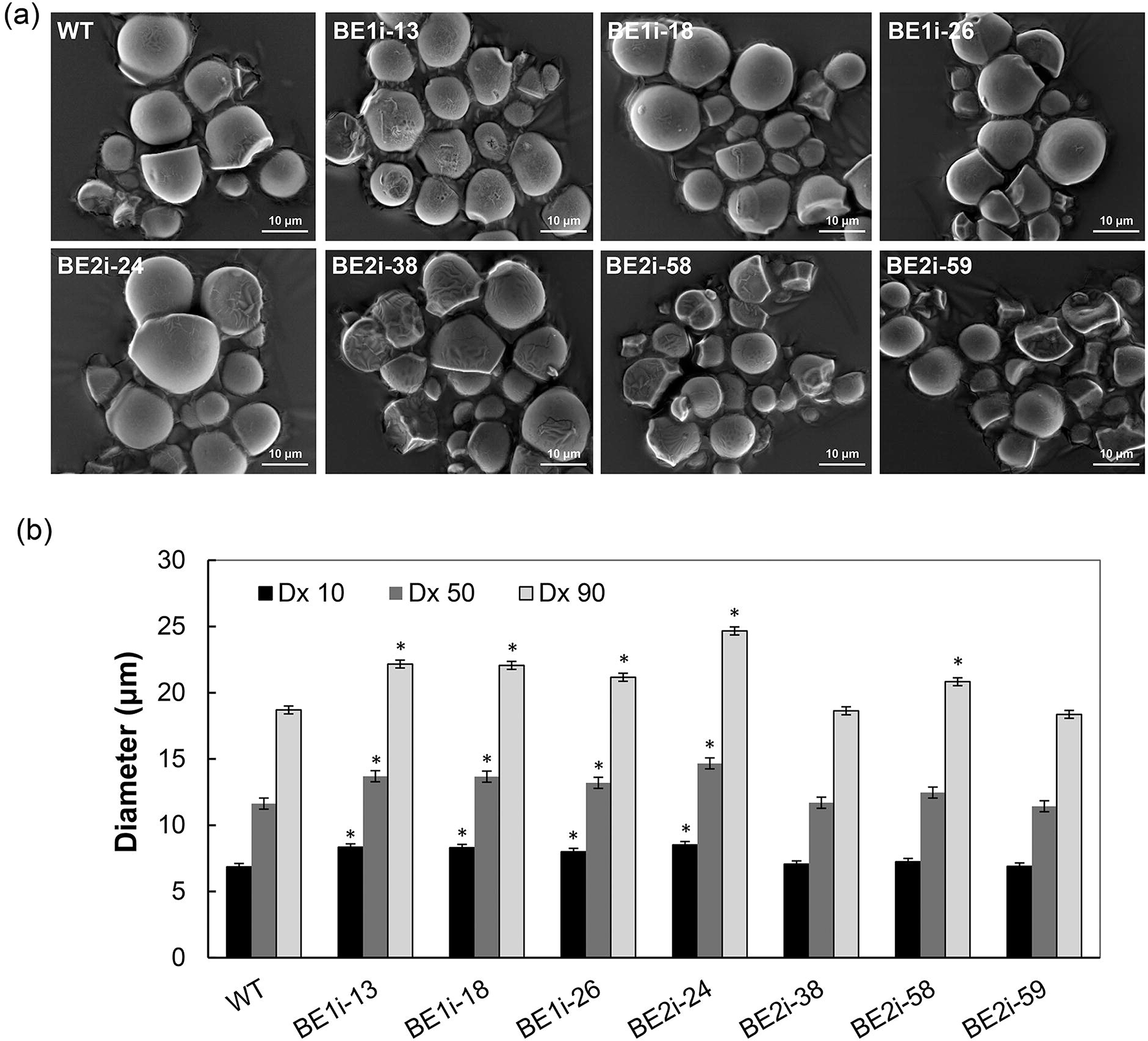
Starch granule morphology from cassava storage roots. Scanning electron microscopy (a) and starch granule size distribution (b) of extracted starches from wild type (WT) and BE-RNAi transgenic lines. BE1i-x, BE1i transgenic plant lines; BE2i-x, BE2i transgenic plant lines. Dx10, Dx50, and Dx90, the projected equivalent diameter below which 10%, 50% and 90% of the total volume of all particles analyzed is represented. Significance was determined by the Student’s *t*-test at * *P* < 0.05.

### MeBE2 rather than MeBE1 leads to dramatic changes in pasting property in cassava storage starch

The BE1i starches exhibited similar pasting patterns to that of WT starch, while the BE2i starches exhibited different patterns (Fig. 5a). More specifically, all three BE1i starches showed increased peak viscosity (PV), hot viscosity (HV), breakdown (BD), and cold viscosity (CV), while decreasing in peak time and pasting temperature (PT) compared to WT starch. Since the value of parameter setback (SB) was correlated with amylose content (Zhou et al., 2015), BE1i starches displayed similar SB values commensurate with the similar amylose content in WT starch (Fig. 3). Meanwhile, due to the increase in amylose content in BE2i-38, BE2i-58, and BE2i-59, starch in these lines showed increased PV, HV, CV, PT, peak time, and SB, but a decrease in BD. These findings indicate that high amylose cassava starch might also be easier to be retrograde than that of WT. HV values represent the resistance to shear thinning. Starch from three BE1i lines (BE1i-13, BE1i-18, BE1i-26) and three BE2i lines (BE2i-38, BE2i-58, BE2i-59) all showed an increased HV over WT. This is especially true for BE2i starches, which ranged from 1129 cP for BE2i-59 to 1136 cP for BE2i-38, which is roughly 2 times higher than that of WT starch (642 cP), suggesting that high amylose cassava starch is superior in terms of resistance to shear thinning. The values of peak time (ranging from 5.5 min for BE2i-59 to 7.0 min for BE2i-58) and PT (ranging from 71.0°C for BE2i-59 to 77.3°C for BE2i-58) in the three BE2i transgenic starches (BE2i-38, BE2i-58, and BE2i-59) were all higher than that of WT starch (5.0 min and 68.9°C, respectively), which indicated that their gelatinization process was slower (Fig. 5a). In addition, the distinctive pasting profile of BE2i-24 starch indicated that its hardness to paste under given conditions renders this line unsuitable for comparison with other starches. However, the severe delay in the gelatinization process suggests the high amylose content may intensively alterthe BE2i-24 starch structure (Fig. 5a).

**Figure 5.**
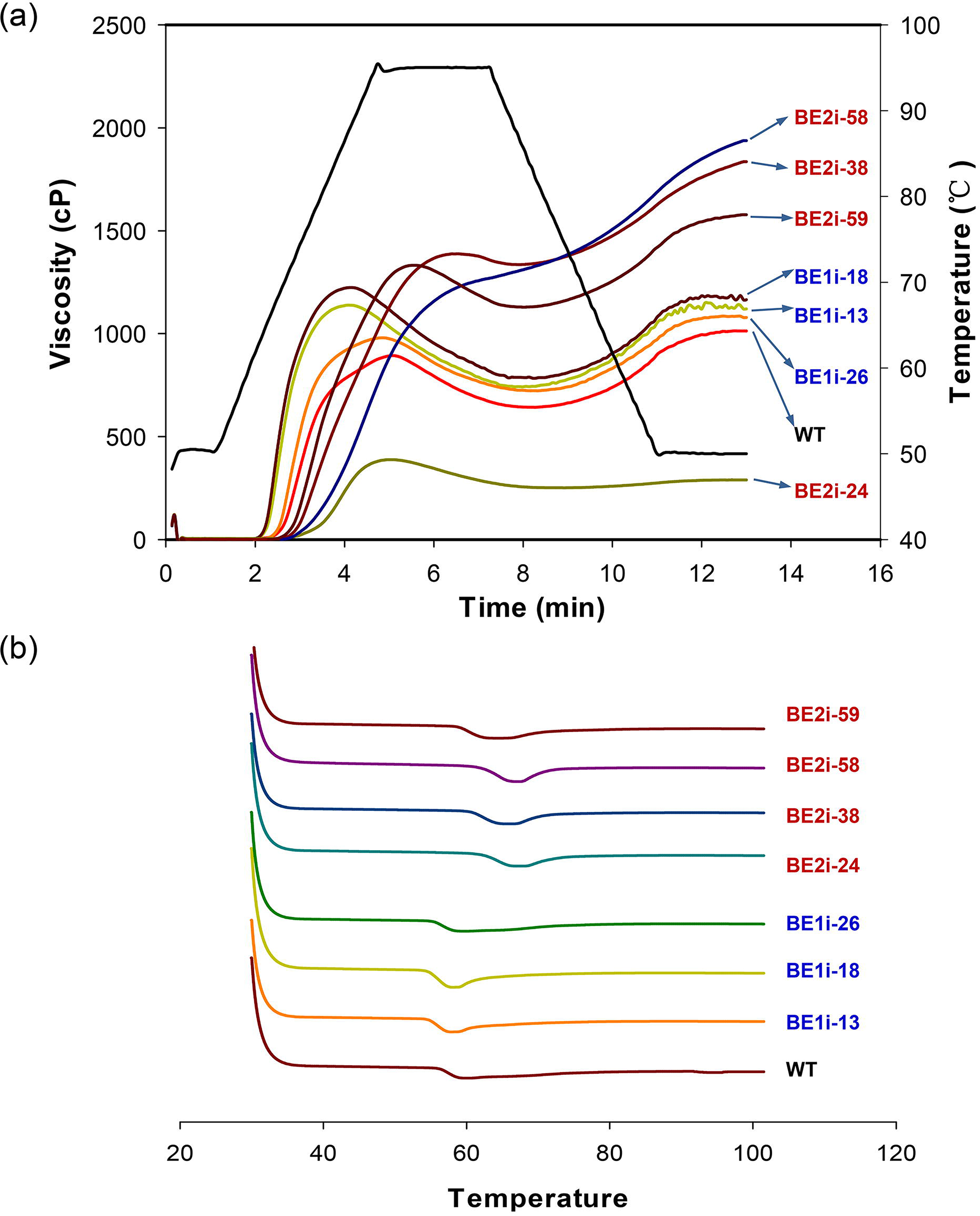
Pasting and thermal properties of cassava storage starches. (a) Rapid Visco-Analyser (RVA) pasting profiles of starches (6% w/v suspension) extracted from storage roots of wild type (WT) and BE-RNAi transgenic lines. (b) Differential scanning calorimeter thermograms of storage starches from WT and BE-RNAi transgenic lines. BE1i-x, BE1i transgenic plant lines; BE2i-x, BE2i transgenic plant lines.

Thermal property analysis of storage starches showed that the melting process requires higher temperature in BE2i starches and lower temperature in BE1i starches compared to WT (Fig. 5b). For the WT starch, the onset temperature (To) and the top melting temperature (Tp) were 56.21°C and 59.34°C, respectively. Both parameters were slightly decreased for all three BE1i starches (T_o_/T_p_: BE1i-13 54.78/57.54°C, BE1i-18 54.58/57.64°C, and BE1i-26 55.34/58.74°C, respectively) and increased for all BE2i starches (T_o_/T_p_: BE2i-24 62.19/66.41°C, BE2i-38 60.61/65°C, BE2i-58 61.94/66.19°C, and BE2i-59 58.84/62.85°C, respectively), especially BE2i-24. The conclusion temperature (Tc) and the thermal (⊿H) were 73.19°C and 12.70 J/g for WT, and their values were decreased in all transgenic starches, especially for BE1i-13 and BE2i-18 starches. The increased To and Tp and decreased Tc indicated that the melting process of the BE2i transgenic starches began later and finished earlier than that of WT, thus showing a lower energy requirement for BE2i starches during the melting process.

### MeBE2 function affects the semi-crystallinity property of storage starch via modified amylopectin structure

X-ray diffraction (XRD) analysis of normal cassava storage starch showed typical A-type crystals with one doublet around 17° (2θ) and 2 singlets predominantly around 15° and 23° (2θ) (Defloor et al., 1998, Zhao et al., 2011). Compared to WT storage starch, BE1i storage starches did not distinctly change in crystallinity pattern (Fig. 6a), whereas all BE2i starches underwent pronounced changes in their diffraction grams, including two new singlets emerging around 5°(2θ) and 20 °(2θ), replacement of a doublet with a singlet appearing around 17 ° (2θ), and replacement of a singlet around 23° (2θ) by one doublet. The peak intensities around 15° (2θ) of BE2i starches were weaker than that of WT and BE1i starches. These XRD features indicated that BE2i storage starches display patterns typical of B-type crystallinity.

**Figure 6.**
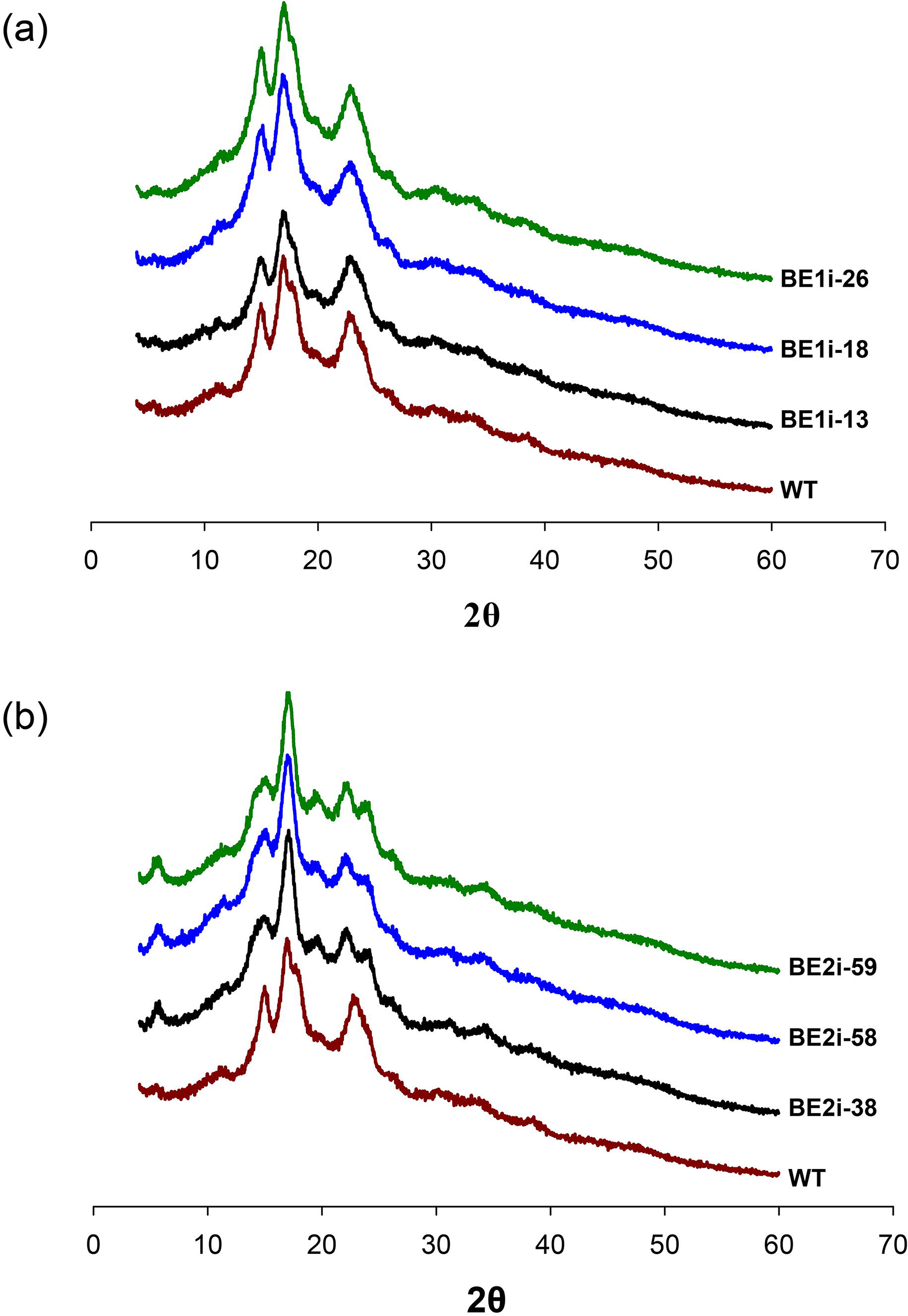
X-ray diffractograms of storage starches from the wild-type (WT) and BE-RNAi transgenic lines. BE1i-x, BE1i transgenic plant lines; BE2i-x, BE2i transgenic plant lines.

Alteration of starch crystal structure and properties in these transgenic starches suggested changes to their amylopectin structure. Chain length distribution analysis of amylopectin by HAPEC-PAD showed differences in the degree of polymerization (DP) between the amylopectins of BE1i or BE2i and WT (Fig. 7). Compared to WT, all three BE1i amylopectins showed a slight increase for long chains of DP >36, and the extent of the difference was less than 0.2% (Fig. 7a). Amylopectins from all three BE2i starches (BE2i-38, BE2i-58, BE2i-59) demonstrated an increase in glucan chains around DP 10-20 and DP >40, but a slight decrease around DP 22-38, with the extent of the difference reaching about 0.5% in BE2i-58 and BE2i-59 (Fig. 7b). The changes in BE2i starches reflect the key function of cassava BE2 during amylopectin biosynthesis.

**Figure 7.**
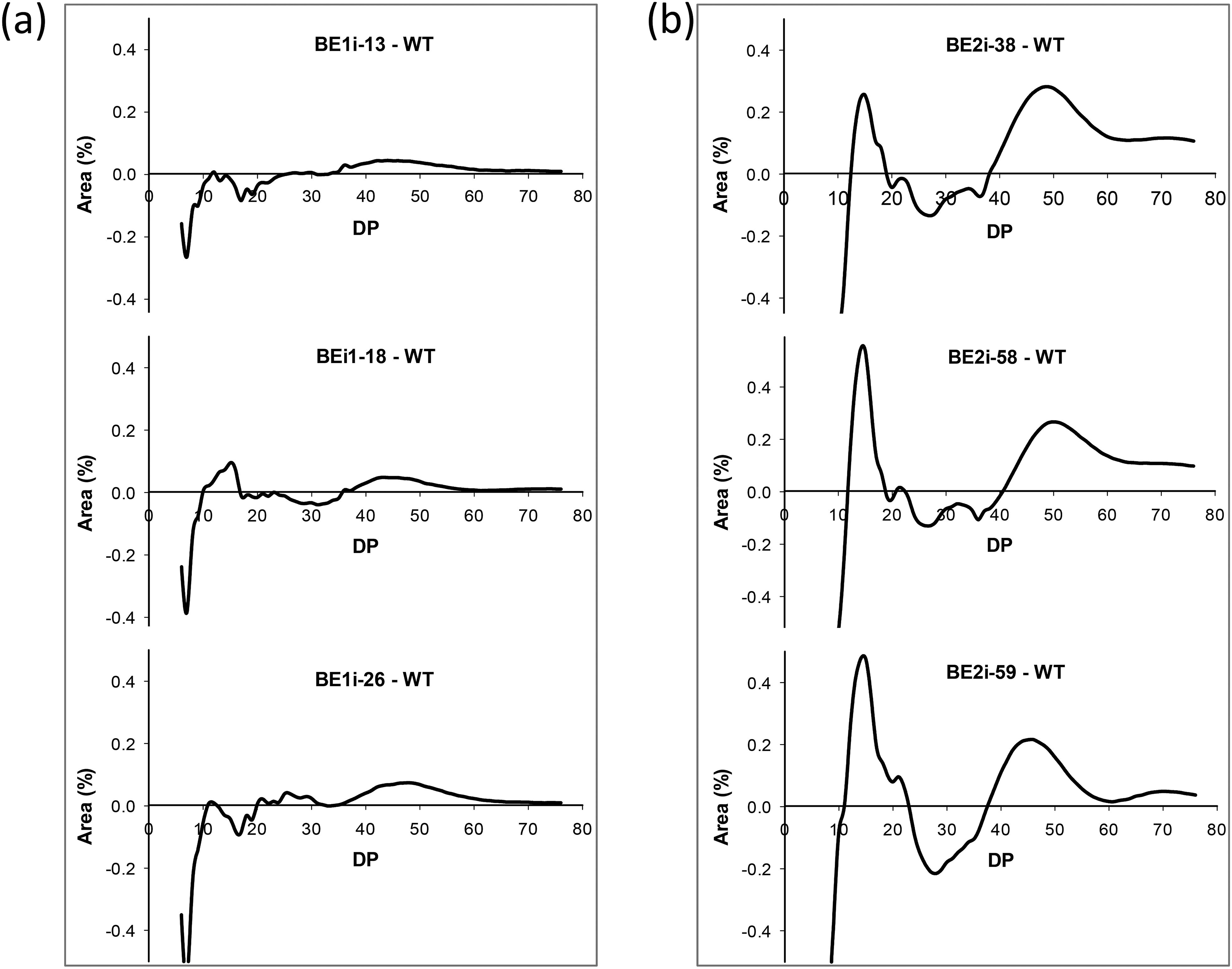
Comparison of chain length distribution of storage starches from wild type (WT) and BE-RNAi transgenic plants. BE1i-x, BE1i transgenic plant lines; BE2i-x, BE2i transgenic plant lines.

## Discussion

High amylose content in starch is a desirable trait in starchy crop breeding and a key characteristic of industrial applications for starch (Jobling 2004). Unlike grains, it is difficult to obtain starch mutants by artificial mutagenesis in root crops like cassava due to their propensity for vegetative propagation, self-pollination, incompatibility, and heterozygosity (Ceballos et al., 2004). Only recently, natural waxy and small granule starch cassava lines were identified from self-pollinated cassava lines in the CIAT cassava breeding program (Ceballos et al., 2007, 2008, 2017). Instead, successful approaches using transgenesis to obtain waxy cassava have been reported in the last decade by down-regulating expression of the *GBSSI* gene either constitutively or tissue-specifically (Raemakers et al., 2005; Zhao et al., 2011; Koehorst-van Putten et al., 2012), or by deletion mutating *GBSSI* and *PTST1* via CRISPR/Cas9-based mutagenesis (Bull et al., 2018). Very-high-amylose cassava starch has not yet been identified through current traditional breeding programs. This need therefore provides an opportunity to generate novel cassava germplasm with high-amylose starch content in storage roots by down-regulating the BE genes that are responsible for amylopectin biosynthesis. In this report, we successfully generated transgenic cassava with very-high-amylose content up to 50% by post-transcriptional silencing of MeBE2, but not MeBE1. Starch physico-chemical analyses confirmed the drastic alteration of starch properties predominately in BE2i starches, indicating the importance of MeBE2 in amylopectin biosynthesis in cassava storage roots.

In higher plants, the starch branching enzymes can be classified into SBEI and SBEII subfamilies which play different roles in amylopectin biosynthesis through distinct differences in their polymer substrate preferences (Tetlow and Emes, 2014). In cassava, the exact functions of BE1 and BE2 are still unclear, although their tissue and spatial expression patterns have been studied under different growth and environmental conditions (Salehuzzaman et al., 1992; Bauama et al., 2003; Pei et al., 2015), or in response to day/night cycle (Supplementary Fig. S1). Our study showed that although down-regulation of MeBE1 or MeBE2 expression in cassava exhibits similar remarkable phenotypic changes and altered circadian oscillation of sugars (Supplementary Fig. S2), only BE2i starches are high-amylose of the B-type, with changes in pasting properties and chain lengths. This finding suggests a fundamental role for BE2 in amylopectin synthesis in cassava storage roots through introduction of more branching points to the starch polymer. Our results are essentially in agreement with research on BE2 function in maize, wheat, rice, pea, and potato (Tetlow and Emes, 2014; Wang et al., 2017).

Silencing MeBE1 expression in cassava has no effect on the amylose content of storage roots, indicating a different role for MeBE1 in cassava starch biosynthesis. Overall, the BE1i starch did not have dramatic changes in starch physico-chemical properties compared to WT except for enlargement of the starch granule size; a similar observation of no significant effects on starch has been reported in maize lacking SBE1 activity (Blauth et al., 2002). Rydberg et al. (2001) reported that potato BE2 was more active than BE1 on an amylopectin substrate, whereas BE1 was more active than BE2 on an amylose substrate. Further investigation is required to decipher the enzymatic process of cassava BE1 protein and its interactions with other starch biosynthetic enzymes (Tetlow et al., 2015). Nevertheless, similar reduction in biomass for both BE1i and BE2i plants indicated that MeBE1 and MeBE2 are equally important for cassava growth and development.

Amylose content is a dominant factor affecting applied properties of starch (Van Hung et al., 2006; Zhou et al., 2015). Important and distinct changes in high-amylose starches from BE2i lines include changes in the paste properties, thermograms, amylopectin chain lengths, and crystallinity. High-amylose cassava starches have increased PV, HV, CV, PT, peak time, and SB; however, BD is decreased, indicating a high capacity for resistance to shear thinning and a slower gelatinization process. These starches also need a higher temperature during the melting process. The transformation of crystal features of high-amylose starch from the A-type to the B-type concurrent with an increase of amylose content in cassava showed a similar trend to that of sweet potato (Zhou et al., 2015), showing three significant alterations at 5 ° (2θ), 17° (2θ) and 20 ° (2θ) in XRD analysis (Fig.6).

The chain length distribution can reflect the branching pattern of amylopectin, which is related to the activity of BEs (Tester et al., 2004; Wang et al., 2017). No remarkable difference between BE1i starch and WT starch was detected except for the very short glucan chains around DP<10, indicating BE1 might not be involved in amylopectin branching. Reduced BE2 function in BE2i lines showed similar trends in increasing glucan chains around DP 10-20 and DP40-60, but decreasing DP22-38. The changes reflect BE2 primary involvement in the branching of amylopectin during starch biosynthesis in storage roots. The increase of DP>40 glucan chains in BE2i starches (Fig.7) may cause the formation of more solid, double-helical crystallites in amylopectin molecules, resulting in the increase in the thermal parameters To and Tp for BE2i lines in DSC analysis (Table II), as reported in other high-amylose starches (Jane et al., 1999; Kong et al., 2015; Zhou et al., 2015). More investigations are required to determine the mechanisms of BE1 and BE2 participation in cassava starch biosynthesis.

In conclusion, our study reports the generation of high-amylose cassava by down-regulating expression of the BE2 gene. The high-amylose cassava starch showed altered starch physico-chemical properties, such as pasting, gelatinization properties, and crystallinity. This work can provide a novel source of cassava starch for various industrial applications. The differences in amylose content and chain length distribution of amylopectin also indicated a divergence in BE1 and BE2 functions in cassava plants.

Further molecular investigation of BE1 and BE2 function in cassava starch biosynthesis, such as enzymatic activity and interaction with other proteins, will improve our understanding of the regulatory mechanisms governing starch accumulation in cassava storage roots.

## Supporting information

## Acknowledgments

This work was supported by the grants from the National Natural Science Foundation of China (31871682), the National Key Technology Research and Development Program of China (2015BAD15B01), the Collaborative Innovation Action—Agricultural Science and Technology Innovation Program of Chinese Academy of Agricultural Sciences (CAAS-XTCX2016009) and the Earmarked Fund for China Agriculture Research System (CARS-11-shzp). We thank Chuanzhong Li and Xinyan Liu for assistance in field experiments and Xiaoyan Gao for SEM experiment.

## Author contributions

W.Z. performed most of the experiments and analyzed the data; S.Z. conducted partial experiments and drafted the manuscript; S.H. and X.H conducted the Western blot analysis and partially starch property assay; W.Z., Q.M. and X.L. maintained transgenic cassava; H.W. assisted with material preparation and discussion; J.Y. coordinated and assisted in preparation the manuscript. P.Z. conceived and designed the study, analyzed the data and revised the paper with input from other authors. All authors discussed the results and approved the manuscript.

## Supplementary files

**Supplementary Figure S1** Transcript changes of starch branching enzymes in leaves (a) and storage roots (b) of cassava plants in response to day/night regime. β-actin was used as an internal control for quantitative comparison of gene expression.

**Supplementary Figure S2** Oscillation in sucrose (a), fructose (b) and glucose (c) of wild type and BE-RNAi transgenic cassava plants during the day/night cycles. WT, wild type; BE1i-x, BE1i transgenic plant lines; BE2i-x, BE2i transgenic plant lines.

---

**Table 1.** Viscosity properties of starches (5%) from wild-type (WT) and BE-RNAi transgenic cassava analyzed using a Rapid Visco-Analyser. Standard deviations are given in parentheses. BE1i-x, BE1i transgenic plant lines; BE2i-x, BE2i transgenic plant lines. The values in the same column with different letters differ significantly (*p* < 0.05).

| Cassava line | Peak viscosity (cP) | Hot viscosity (cP) | Break down (cP) | Cold Viscosity (cP) | Setback (cP) | Peak time (min) | Pasting temperature (°C) |
| --- | --- | --- | --- | --- | --- | --- | --- |
| WT | 893 ±7.81f | 642 ±5.86f | 251±6.43c | 1013±13.80f | 371±7.94d | 5.0±0.04 d | 68.9±0.6e |
| BE1i-13 | 1139 ±17.04d | 740±17.21e | 399±2.00b | 1121±34.39de | 381±27.23d | 4.1±0.04f | 63.6±0.03g |
| BE1i-18 | 1225±8.50c | 780±11.02d | 445±10.15a | 1165±21.22d | 386±14.98d | 4.1±0.04f | 63.6±0.08g |
| BE1i-26 | 980±8.96e | 721±9.54e | 259±10.12c | 1079±19.50ef | 358±10.26d | 4.8±0.10e | 67.5±0.48f |
| BE2i-24 | 388 ±9.64g | 251±3.61g | 137±6.08e | 289±4.93g | 38±3.06e | 5.0±0.04de | 81.3 ±0.46a |
| BE2i-38 | 1390±12.42a | 1336±21.66a | 54±11.72g | 1835±31.47b | 499±14.64b | 6.5±0.12b | 72.7±0.05c |
| BE2i-58 | 1258±18.95c | 1169±22.03b | 89±4.58f | 1938±40.50a | 769±19.67a | 7.0±0.00a | 77.3±0.48b |
| BE2i-59 | 1333±8.33b | 1129±2.52c | 204±10.82d | 1578±15.95c | 449±14.11c | 5.5±0.07c | 71.0±0.03d |

**Table 2.** Thermal properties of storage starches from wild-type (WT) and BE-RNAi transgenic lines as determined by differential scanning calorimetry (DSC). Standard deviations are given in parentheses. BE1i-x, BE1i transgenic plant lines; BE2i-x, BE2i transgenic plant lines; To, Onset temperature; Tp, Peak temperature; Tc, End temperature; ΔH, Endothermic enthalpy. The values in the same column with different letters differ significantly (*p* < 0.05)

| Cassava line | T<sub>o</sub> (°C) | T<sub>p</sub> (°C) | T<sub>c</sub> (°C) | ΔH (J/g) |
| --- | --- | --- | --- | --- |
| WT | 56.21±0.11d | 59.34±0.18d | 73.19±0.3a | 12.70±0.55a |
| BE1i-13 | 54.78±0.02f | 57.54±0.04f | 61.06±0.35c | 9.71±0.43b |
| BE1i-18 | 54.58±0.02f | 57.64±0.03f | 61.31±0.43c | 9.68±0.59b |
| BE1i-26 | 55.34±0.1e | 58.74±0.14e | 71.12±0.36b | 12.42±0.52a |
| BE2i-24 | 62.19±0.27a | 66.41±0.05a | 72.33±0.49a | 11.47±0.92ab |
| BE2i-38 | 60.61±0.06b | 65±0.02b | 70.49±0.14b | 12.70±0.59a |
| BE2i-58 | 61.94±0.1a | 66.19±0.2a | 71.12±0.34b | 12.59±0.08a |
| BE2i-59 | 58.84±0.24c | 62.85±0.09c | 70.41±0.23b | 11.67±1.05a |

