## Supplementary figures and images for "Production of very-high-amylose cassava by post-transcriptional silencing of branching enzyme genes"

### Supplementary file 1

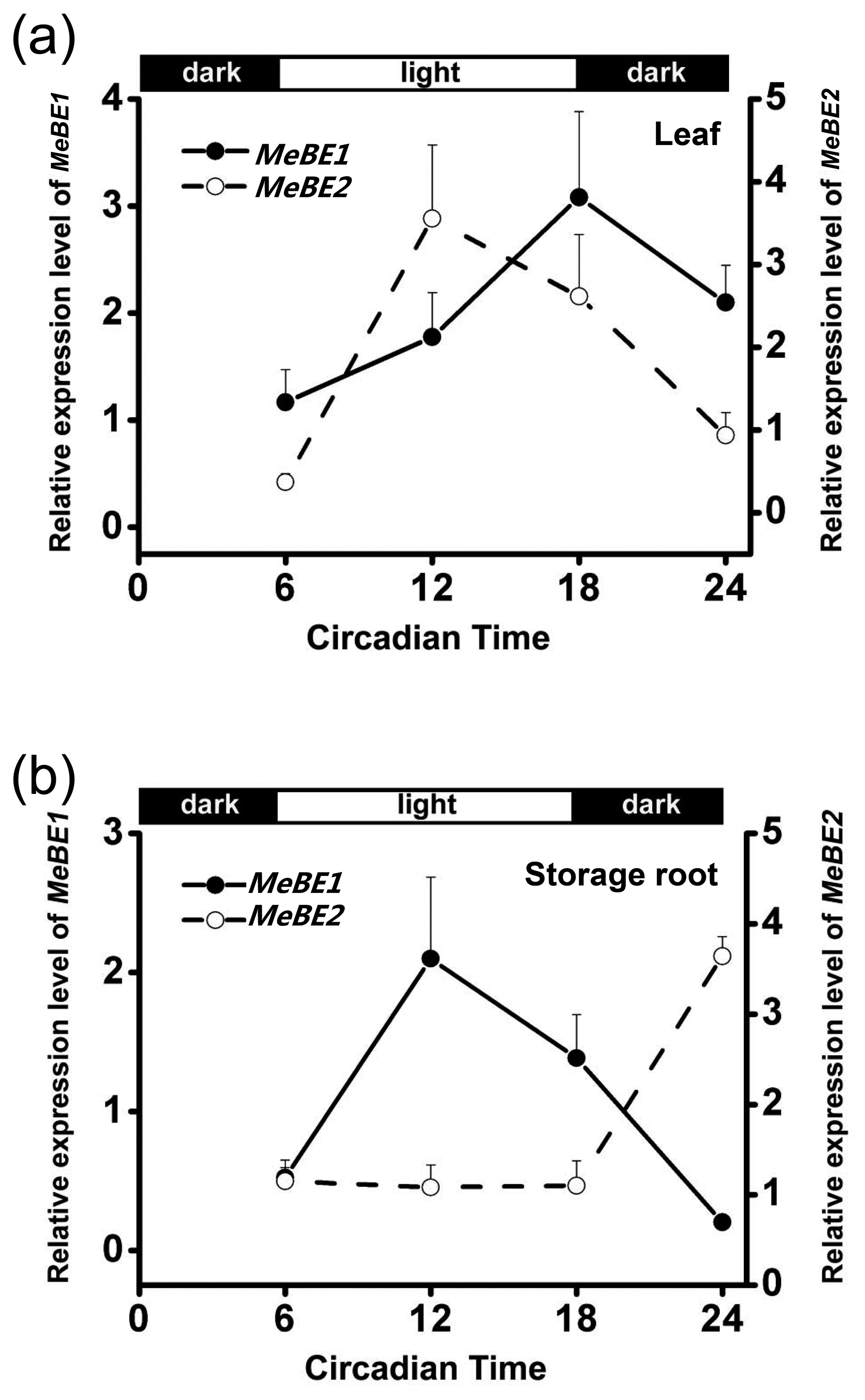

### Supplementary file 2

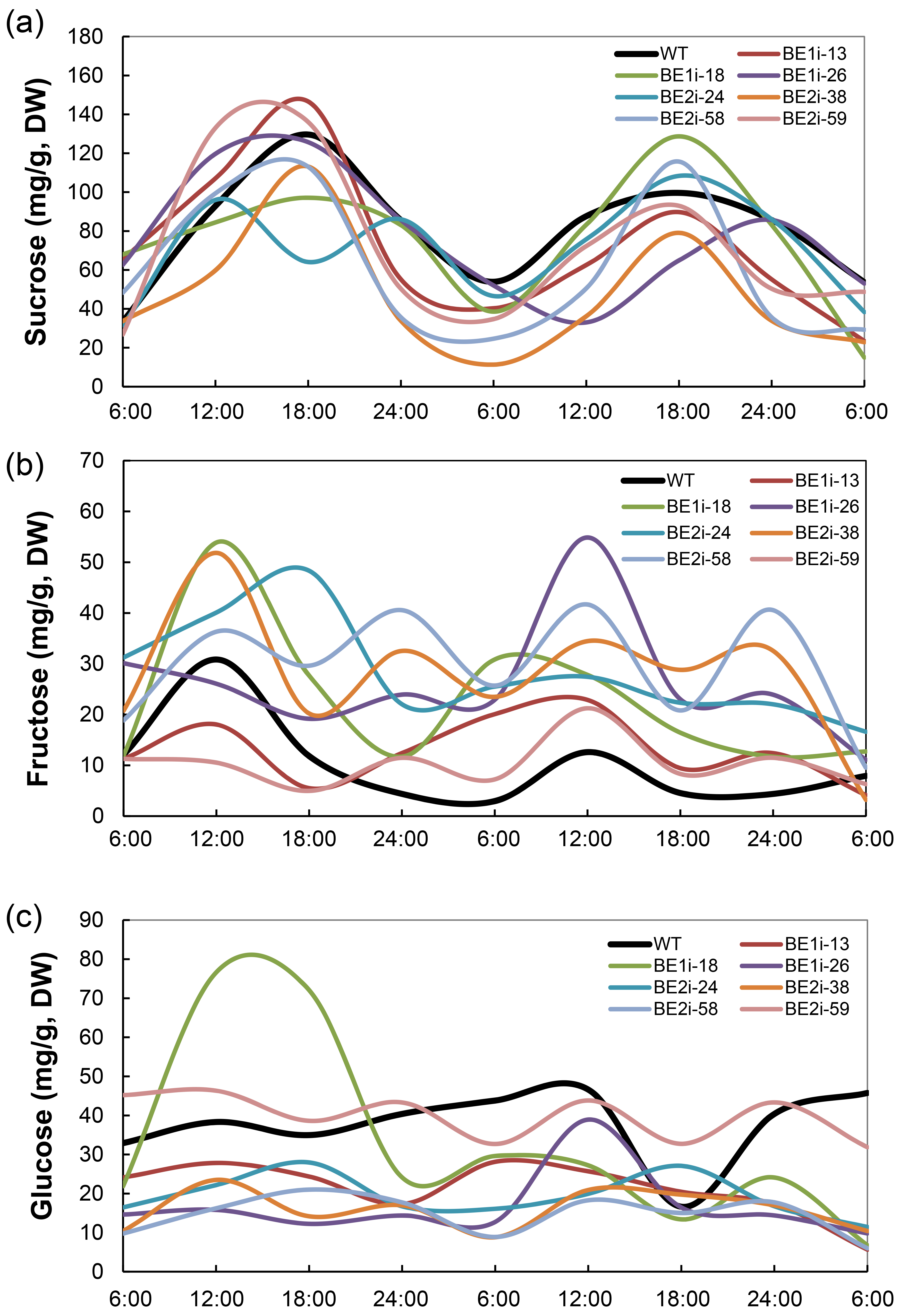
